# SPEAR: Predicting Gene Expression from Single-Cell Chromatin Accessibility

**DOI:** 10.64898/2026.04.13.717809

**Authors:** Thussenthan Walter-Angelo, Yasin Uzun

## Abstract

Single-cell multiome assays enable direct measurement of chromatin accessibility and gene expression within the same cell. Still, most experimental designs remain constrained to two (and, less commonly, three) modalities per cell. This limitation motivates computational models that can predict unmeasured layers and, simultaneously, help dissect how cis-regulatory accessibility relates to transcription at gene resolution. Existing cross-modal methods often prioritize latent alignment or modality reconstruction, making it difficult to isolate the impact of model inductive bias under a shared cis-regulatory feature definition. We present SPEAR, a configuration-driven framework for gene-centric regression of single-cell gene expression from chromatin accessibility using a fixed transcription-startsite–centered representation shared across model families. Here we show that, under identical features, splits, and evaluation, model performance stratifies reproducibly across two multiome systems (mouse embryonic development and human hemogenic endothelium), with transformer encoders achieving the strongest mean test correlations (0.546 and 0.470, respectively). Per-gene performance distributions reveal substantial heterogeneity in predictability, indicating that accessibility-driven signal is concentrated in a subset of genes across contexts. Shapley value–based feature attribution further localizes predictive signal to promoter-proximal bins, with feature importance decaying with distance from the transcription start site, supporting a promoter-centered regime of cis-regulatory control within the modeled window. Together, these results provide a controlled comparison of inductive biases for chromatin-to-expression prediction and deliver analysis-ready outputs for gene-level interpretation.

SPEAR is open source and publicaly available for use at https://github.com/UzunLab/SPEAR.

Supplementary data are available.

## Introduction

Single-cell genomics has transformed our ability to study gene regulation by resolving cell-to-cell heterogeneity that is obscured in bulk measurements. Early single-cell RNA sequencing (scRNA-seq) established that transcriptional states can be profiled at cellular resolution and used to define cell types, trajectories, and regulatory programs (1, 2). In parallel, single-cell chromatin accessibility assays (scATAC-seq) enabled the measurement of regulatory DNA activity, providing a complementary view of the epigenetic landscape underlying transcriptional potential (3). Together, these technologies motivated a central question in gene regulation: to what extent can gene expression be predicted from chromatin accessibility, and what does that predictability reveal about cis-regulatory control? A significant step forward came from paired multimodal assays that capture two (or more) modalities in the same cell, enabling direct alignment between chromatin state and transcriptional output. Early paired assays demonstrated joint profiling of accessibility and RNA in thousands of single cells, allowing quantification of chromatin–expression coupling without post hoc integration across separate experiments (4). More recently, high-throughput multiome platforms have made joint scATAC+scRNA measurements broadly accessible, accelerating development of computational methods for cross-modality mapping and prediction (4, 5). A growing ecosystem of methods addresses cross-modality prediction, but they often differ in objective and interpretability. Some approaches learn shared latent representations and translate between modalities, treating RNA prediction primarily as a reconstruction task. BABEL is a canonical example that demonstrates ATAC-to-RNA translation after training on paired data (6). More recent generative and integration frameworks similarly enable cross-modal imputation or alignment, often prioritizing shared latent representations and modality reconstruction over explicit gene-centric regression diagnostics (7, 8). In contrast, other methods focus on gene-level regulatory modeling and enhancer interpretation (often with constrained model classes), aiming to learn interpretable cis effects or nominate functional elements rather than benchmark broad architectural choices (9, 10). Still other work uses paired multiome regression to identify functional enhancers and quantify regulatory potential, highlighting how paired prediction can act as a lens into cis-regulatory logic and disease-relevant control (10). However, a gap remains: we lack a controlled, gene-centric benchmarking framework that compares fundamentally different model families under a standard, fixed cis-regulatory representation, producing standardized, reproducible outputs suitable for downstream biological interpretation. Many published comparisons confound model choice with differences in feature construction (peak-to-gene linking, gene activity definitions, windowing schemes), training objectives, or evaluation protocols, making it challenging to attribute performance differences to inductive bias rather than pipeline idiosyncrasies (7). Moreover, even when prediction performance is reported, gene-level heterogeneity (which genes are predictable vs not), split-wise generalization behavior, and feature-attribution structure are often not produced as first-class outputs. Yet, these are precisely the diagnostics needed to connect prediction to regulatory mechanisms. Furthermore, because chromatin–expression coupling and data characteristics vary across biological systems, the optimal model family is not guaranteed to transfer across datasets, motivating systematic benchmarking when applying chromatin-to-expression prediction to a new context. To address this gap, we present SPEAR (Single-cell-based Prediction of Gene Expression from Chromatin Accessibility Readouts), a modular framework for formulating chromatin-to-expression modeling as supervised regression on paired single-cell multiome data. SPEAR uses a deterministic, gene-centric cisregulatory representation constructed from chromatin accessibility in fixed genomic windows centered on each gene’s transcription start site. By holding the feature definition, data splits, and evaluation protocol constant, SPEAR enables systematic comparisons among diverse model families that differ in inductive bias (e.g., linearity, nonlinearity, locality, sequence structure, attention, and relational structure). In addition to aggregate accuracy measures, the framework supports gene-resolved evaluation and interpretability analyses by producing standardized per-gene performance summaries and feature-attribution outputs aligned to genomic coordinates. SPEAR is configuration-driven and designed to facilitate reproducible experimentation and targeted extensions, such as alternative window sizes, feature selection strategies, and incorporation of additional cis or trans features. In this study, we apply SPEAR to paired scATAC-seq and scRNA-seq datasets spanning distinct biological contexts (mouse embryonic development and human hemogenic endothelium) to characterize how model family and inductive bias interact with a fixed promoter-centered representation.

## Methodology

### Overview

SPEAR (Single-cell-based Prediction of Gene Expression from Chromatin Accessibility Readouts) is a supervised learning framework for predicting gene expression at single-cell resolution from chromatin accessibility. Using paired scATAC-seq and scRNA-seq measurements from multiome assays, SPEAR formulates chromatin-to-expression mapping as a regression problem, predicting normalized RNA expression values from cis-regulatory accessibility features. An overview of the SPEAR workflow is shown in Figure 1.

**Fig. 1.**
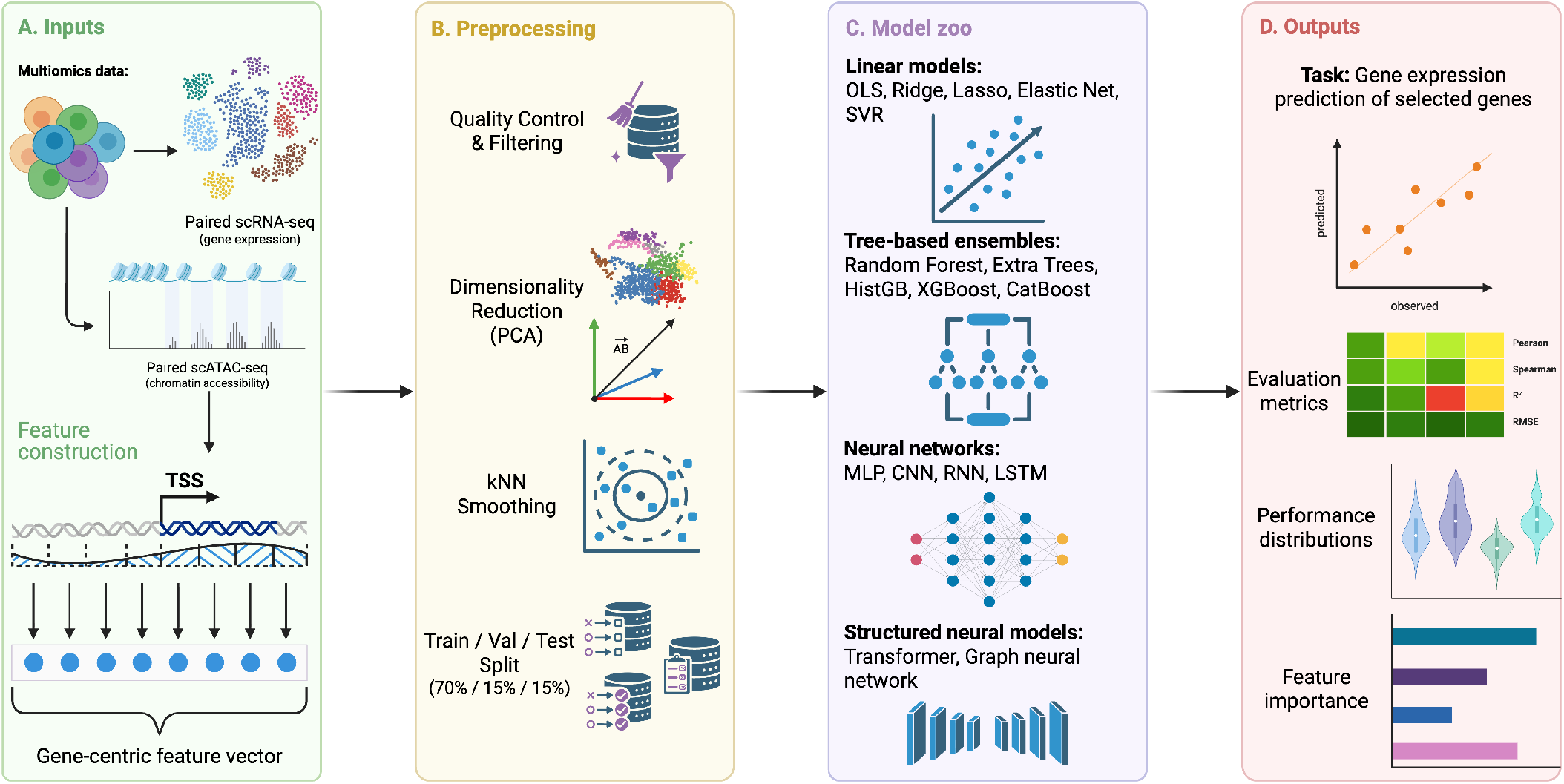
Overview of the SPEAR framework. (A) Paired single-cell chromatin accessibility (scATAC-seq) and gene expression (scRNA-seq) profiles from multiome assays are used as input. Chromatin accessibility is aggregated into gene-centric cis-regulatory features by binning fixed genomic windows centered on transcription start sites (TSS). (B) Preprocessing includes quality control, dimensionality reduction, optional k-nearest neighbor smoothing, and dataset splitting. (C) A modular model zoo spanning linear models, tree ensembles, deep neural networks, and structured architectures is evaluated under an identical feature representation. (D) SPEAR outputs standardized benchmarking artifacts, including predictions, performance summaries, gene-level distributions, and feature-importance profiles for reproducible downstream analysis. *Created with BioRender*.*com*.

Let *X*_*i,g*_ ∈ ℝ^*B*^ denote the cis-regulatory accessibility feature vector for gene *g* in cell *i*, constructed from *B* genomic bins surrounding the transcription start site, and let *y*_*i,g*_ ∈ ℝ denote the corresponding normalized gene expression value. SPEAR formulates chromatin-to-expression prediction as supervised regression:

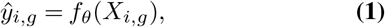

where *f*_*θ*_ is a model parameterized by *θ*, drawn from a specified model family.

In the multi-output setting, features for all target genes are concatenated per cell and the model jointly predicts expression for all genes:

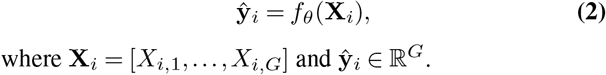

where **X**_*i*_ = [*X*_*i*,1_, …, *X*_*i,G*_] and ŷ_*i*_ ∈ ℝ^*G*^.

### Data preprocessing and normalization

Paired scATAC-seq and scRNA-seq measurements were assumed to originate from the same biological cells. Cell barcodes were intersected and reordered to ensure consistent alignment between accessibility features and expression targets. RNA expression was derived from raw gene-level count matrices and normalized using counts per million (CPM), followed by a log1p transformation (log(CPM+1)). When pre-normalized expression layers were available, redundant normalization was avoided. The resulting matrix served as the regression target for all models. Chromatin accessibility values were computed from ATAC fragment counts and normalized prior to feature aggregation. To stabilize learning with sparse single-cell measurements, all benchmarking results reported here use the default configuration: PCA-informed k-nearest neighbors (kNN) smoothing.

### Gene-centric cis-regulatory feature construction

For each gene, SPEAR constructs a cis-regulatory feature representation from chromatin accessibility within a fixed genomic window centered on the transcription start site (TSS). By default, a ±10 kb window around the annotated TSS was used to capture promoter-proximal regulatory activity. Each window was subdivided into 40 non-overlapping bins of equal width (500 bp). Bins were oriented relative to the gene strand to ensure consistent upstream and downstream positioning. For each cell, ATAC fragments overlapping each bin were aggregated, yielding a fixed-length 40-dimensional feature vector per gene per cell. This representation is independent of gene length, peak density, and total accessibility counts. Feature matrices were deterministically regenerated from fragment files and gene annotations for every run. Gene order, bin order, and feature indexing were tracked and stored alongside model outputs to ensure reproducibility. Depending on model architecture, gene-level feature vectors were either modeled independently or concatenated across genes to form a single high-dimensional input per cell. All runs in this study use SPEAR to train multi-output models that jointly predict expression for all genes in the provided gene list.

### Model families

SPEAR benchmarks a diverse set of regression models chosen to represent distinct inductive biases for chromatin-to-expression prediction. All models operate on the same gene-centric, TSS-centered cis-regulatory feature representation and are trained to predict normalized gene expression from chromatin accessibility, enabling controlled comparisons across architectures. The model zoo includes linear regressors (OLS, ridge, lasso, elastic net), tree-based ensembles (random forest, Extra Trees, XGBoost, CatBoost), and neural architectures spanning multilayer perceptrons (MLPs), sequence-structured models (CNNs, RNNs/LSTMs, transformer encoders), and graph neural networks (GNNs). Linear and tree-based models provide interpretable and nonlinear baselines, while neural models capture increasingly structured dependencies across ordered cis-regulatory bins, from local (CNNs) and directional (RNNs) to global attention-based interactions (transformers). GNNs evaluate the utility of explicitly relational inductive biases. All model architectures and hyperparameters were defined via external configuration files and logged for reproducibility.

### Training protocol and evaluation

All models were trained using identical train, validation, and test splits within each dataset. When group annotations were available, group-aware splitting was used to prevent information leakage; otherwise, random splitting was applied. Neural network models were trained using gradient-based optimization with early stopping based on validation loss. The best-performing checkpoint was retained for test evaluation. Random seeds were fixed where applicable to reduce stochastic variability. Model performance was assessed on held-out test cells using multiple complementary metrics. The primary evaluation was based on Pearson correlations between predicted and observed gene expression, computed per gene and summarized across genes. Additional metrics included root mean squared error (RMSE), Spearman correlation, and the coefficient of determination (*R*^2^). For each run, SPEAR produced standardized outputs including raw test-set predictions, per-gene performance summaries, predicted-versus-observed scatter plots, and training diagnostics. All outputs, along with configuration metadata, were persisted to disk to support reproducibility and downstream analysis.

## Experimental setup

### Datasets and biological systems

SPEAR was evaluated on paired single-cell chromatin accessibility and gene expression data from two biological systems: (i) a mouse embryonic multiome dataset and (ii) a human hemogenic endothelium–derived multiome dataset.

### Mouse embryonic multiome

The embryonic dataset was obtained from GEO accession GSE205117 (11). This study profiled early mouse organogenesis using the 10x Genomics Multiome assay, generating paired snRNA-seq and snATAC-seq measurements across a developmental time course spanning embryonic day E7.5 to E8.75. The original dataset includes wild-type replicates and a CRISPR-mediated Brachyury (T) knockout condition. For this study, only wild-type cells were retained, and all CRISPR perturbation samples were excluded to avoid confounding regulatory effects unrelated to baseline chromatin–expression coupling.

### Human hemogenic endothelium–derived multiome

The endothelial dataset was obtained from GEO accession GSE270141 (12). This dataset consists of paired scRNA-seq and scATAC-seq profiles generated using the 10x Genomics Multiome platform from hemogenic endothelium–derived populations cultured under hypoxic (4% O_2_) or normoxic conditions. From the whole dataset, only endothelial cells were retained for analysis, while non-endothelial populations present in the original study were filtered out to maintain a homogeneous cell population.

Across both datasets, RNA and ATAC measurements were treated as originating from the same individual cells. Cell barcodes were intersected and aligned across modalities before feature construction to ensure correct pairing of chromatin features and expression targets.

### Reference annotation files and genome resources

Gene coordinates and transcription start sites (TSS) were defined using reference gene annotations appropriate to each organism. For the mouse embryonic dataset, gene models were derived from the NCBI RefSeq annotation corresponding to the GRCm39 reference genome (GCF_000001635.27), which provides curated gene structures and TSS definitions for Mus musculus (13). For the human endothelial dataset, annotations were derived from GENCODE v44, a comprehensive gene annotation for the GRCh38 human reference genome that integrates evidence from multiple transcript sources (14). These annotations were used consistently throughout feature construction to define TSS-centered cisregulatory windows, ensuring reproducible gene definitions and stable mapping between chromatin accessibility features and gene expression targets across runs.

### Dataset preprocessing and quality control

For both datasets, preprocessing produced paired, QC-filtered AnnData objects (.h5ad) that served as the direct inputs to the SPEAR pipeline.

### Mouse embryonic dataset preprocessing

Raw persample RNA matrices and a shared ATAC peak matrix were processed on a per-sample basis. RNA and ATAC barcodes were harmonized by renaming RNA barcodes to include sample identifiers and subsetting ATAC columns to the intersecting barcode set. Cells were retained only if present in both modalities. RNA quality control filtered out cells with mitochondrial RNA fractions exceeding 5% (mitochondrial genes defined by MT-prefixes). Low-complexity filtering was applied to both RNA and ATAC matrices, requiring a minimum of 200 detected genes per cell and at least 3 cells per gene. Per-sample QC-filtered matrices were cached and subsequently merged across samples to produce combined RNA and ATAC matrices of 54,301 paired cells. Sample identifiers were retained in cell metadata and later used for group-aware data splitting.

### Human endothelial dataset preprocessing

Raw RNA and ATAC data were provided as R SingleCellExperiment objects and converted to AnnData format. Gene symbols were attached using a reference RNA-only mapping, and RNA and ATAC modalities were aligned by intersecting barcodes. RNA QC filtering removed cells with mitochondrial RNA fractions exceeding 15%, a threshold selected to retain approximately 5,000 high-quality endothelial cells. As in the embryonic dataset, low-complexity filtering required at least 200 detected genes per cell and at least 3 cells per gene, and only cells retained in both modalities after QC were kept. The final processed inputs consisted of paired RNA and ATAC AnnData objects containing only 4,735 paired endothelial cells. Across both datasets, all preprocessing steps were applied before model fitting and held fixed across model families to avoid confounding effects from modality-specific filtering.

### Gene selection, feature dimensionality, and fallback manifests

To balance biological coverage with computational feasibility, experiments were scoped to a fixed target set of 1,000 genes per dataset. Gene selection was performed randomly from an eligible gene pool, with expression-based filtering applied to remove extremely sparse targets. Specifically, candidate genes were required to be expressed (expression ≥0) in at least 10% of cells, with a minimum number of expressing cells enforced during dataset construction. Random sampling was performed without replacement using a fixed random seed to ensure reproducibility, and the resulting gene lists were stored as gene manifest files reused across all model families. For each gene, cis-regulatory features comprised 40 bins spanning ±10 kb around the TSS, yielding 40,000 input features per cell when modeling 1,000 genes jointly. This extremely high-dimensional input space poses a substantial challenge for certain model classes, particularly linear and margin-based models, where optimization over tens of thousands of correlated features can be unstable or prohibitively slow. In practice, some models failed to make meaningful optimization progress on the complete 1,000-gene task within reasonable time limits. To ensure robust benchmarking coverage, a 100-gene fallback manifest was maintained, comprising genes that reliably passed all preprocessing and training checks across model families. This fallback was used for debugging, rapid iteration, and for models that could not feasibly converge at the 1,000-gene scale. Notably, support vector regression failed to train even on the 100-gene embryonic dataset, highlighting the practical limitations of specific classical methods in this regime.

### Model execution and controlled benchmarking

All models were trained and evaluated using identical gene manifests, data splits, and cis-regulatory feature representations within each dataset. Where applicable, group-aware train/validation/test splits were used to prevent information leakage across biological replicates; otherwise, standard random splits were applied. Models were executed using computational resources appropriate to their implementation (e.g., GPU acceleration for deep learning models and CPU execution for classical regressors). Still, resource allocation did not affect data inputs or evaluation procedures. Large benchmarking sweeps were executed in parallel using a job scheduler on a high-performance computing environment. Importantly, all comparisons reported in this study reflect differences in model inductive bias and optimization behavior rather than differences in preprocessing, feature construction, or evaluation protocol.

## Results

### Predictive performance varies systematically across model families and biological contexts

The main design principle of SPEAR is the use of a fixed, gene-centric feature representation shared across all model families. This concept enables controlled, systematic comparison of diverse regression approaches under identical inputs, training procedures, and evaluation criteria. We first evaluated predictive performance across model families on held-out test cells in the two datasets (Figure 2). Performance was quantified using Pearson correlation between predicted and observed gene expression, computed per gene and summarized across genes. Across both datasets, models showed apparent, reproducible differences in test accuracy (Figure 2A).

**Fig. 2.**
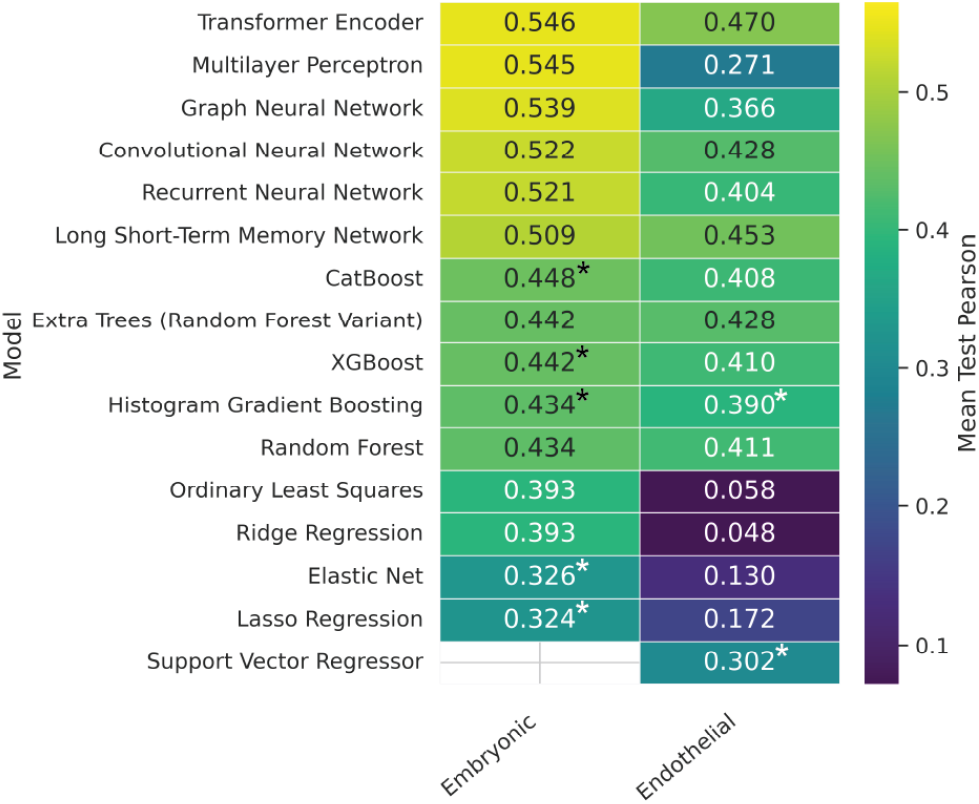
Model performance across biological contexts. Heatmap of mean test-set Pearson correlations (computed per gene and summarized across genes) for each model family in the embryonic and endothelial datasets under an identical promoter-centered representation.

Despite significant differences in organism and experimental context, the highest-performing methods were consistently deep neural architectures, with the transformer encoder ranking first in both datasets. In the embryonic dataset, the transformer encoder achieved the top performance (mean test correlation of 0.546), closely followed by the MLP (0.545) and GNN (0.539). In the endothelial dataset, the transformer encoder was the top performer (0.470), followed by LSTM (0.453) and CNN (0.428). In contrast, classical baselines were substantially weaker, particularly in the endothelial dataset. For example, the correlation for ridge and ordinary least squares (OLS) regression was close to zero (0.048 and 0.058, respectively). Tree ensembles showed intermediate accuracy, with correlation values higher than those of statistical regression but lower than those of deep learning (random forest: 0.434 in the embryonic dataset and 0.411 in the endothelial dataset; Cat-Boost: 0.448 and 0.408 for the same datasets). While the top-performing model was consistent across datasets, the relative ordering among several strong alternatives differed between systems, and absolute performance also shifted (Figure 2B). Beyond the transformer encoder, model rankings diverged in a dataset-dependent manner: the embryonic dataset favored a generic nonlinear baseline (MLP) and a relational model (GNN) that performed comparably to the transformer, whereas the endothelial dataset favored sequence-structured models (LSTM and CNN). Many models also attained higher correlations in the embryonic setting (e.g., transformer encoder 0.546) than in the endothelial setting (0.470), suggesting that dataset-specific biological and technical factors modulate how strongly local cis-accessibility predicts expression. Together, these shifts indicate that no single “runner-up” architecture is universally optimal, motivating systematic benchmarking of new datasets across model families to identify inductive biases best matched to the underlying regulatory context: an application directly supported by SPEAR.

### Gene-level predictability exhibits broad heterogeneity across models and datasets

To move beyond model-level averages, we examined the distribution of per-gene predictive performance across model families (Figure 3). Across both datasets, per-gene Pearson correlations showed broad distributions, indicating that chromatin-to-expression predictability is strongly gene dependent.

**Fig. 3.**
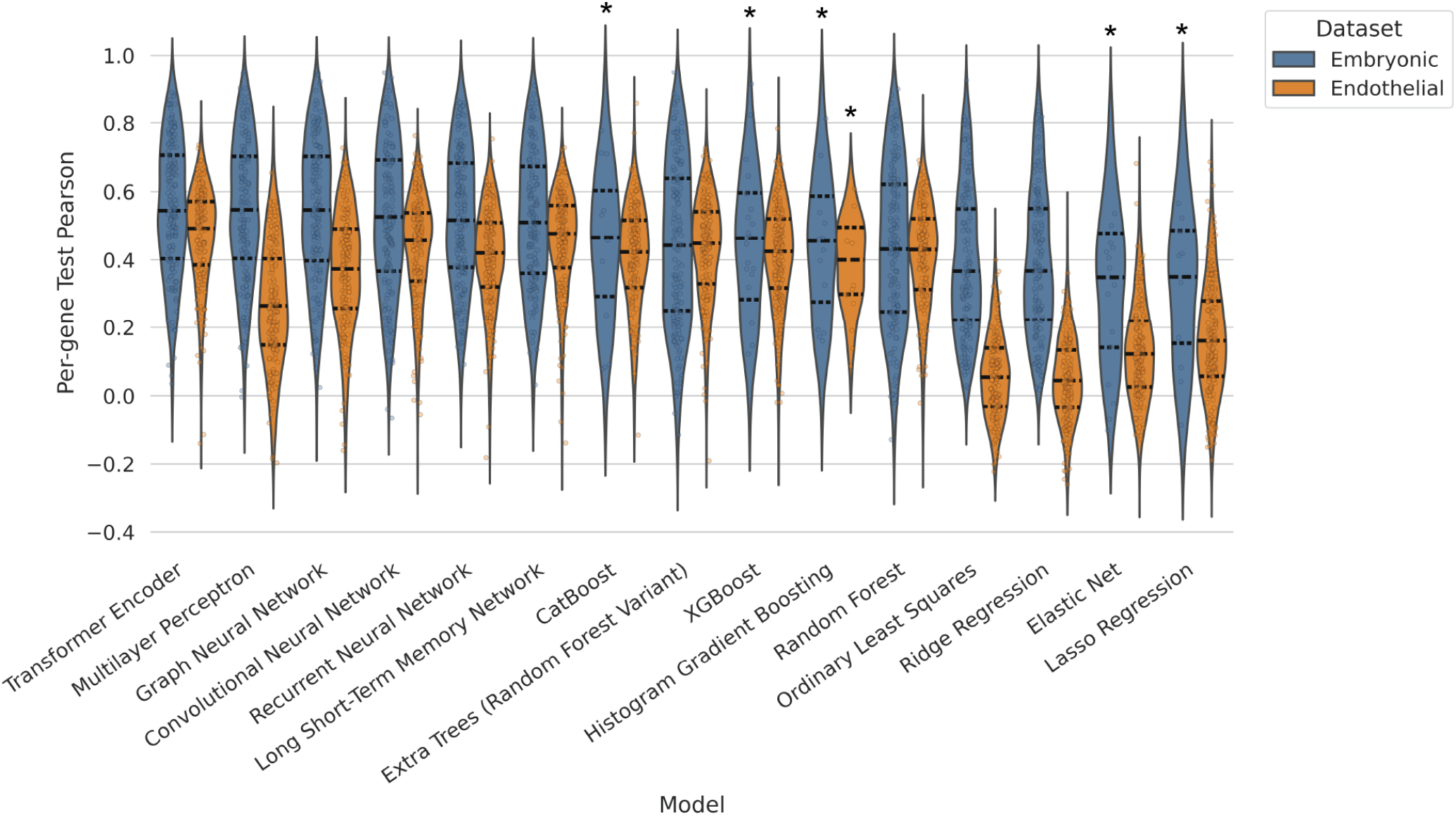
Overall predictive performance across datasets. Violin plots showing the distribution of per-gene test-set Pearson correlations between predicted and observed expression for each model family, stratified by dataset. Distributions emphasize gene-level heterogeneity in predictability beyond mean performance.

Deep models improved not only average performance but also the upper tail of predictability. This result is consistent with the idea that higher-capacity sequence/structure-aware models are better able to exploit subtle but reproducible cisregulatory patterns (e.g., strand-consistent promoter asymmetry and local bin interactions) that may be diluted under linear assumptions. At the same time, the persistence of low-performing genes even under the transformer encoder suggests that many genes exhibit expression variation driven by factors not captured by the local accessibility window (e.g., distal regulation, TF abundance, cell-state effects, or technical dropout). Notably, distributions are shifted upward in the embryonic dataset relative to the endothelial dataset, consistent with higher mean test performance in embryonic across top models (e.g., for the transformer encoder, 0.546 in embryonic vs 0.470 in endothelial). This fact supports the interpretation that the strength and locality of cis-regulatory signals differ across biological contexts.

### Models generalize consistently across data splits, with overfitting concentrated in classical ensembles

We next compared performance across train/validation/test splits (Figure S1) and quantified generalization gaps (Figure 4). For the best-performing deep models, the train-to-test gap was modest, suggesting that improvements reflect the extraction of real signal rather than memorization. In the embryonic dataset, the transformer encoder showed the smallest generalization gap (mean correlation of 0.668 for training and 0.546 for test, with a gap of 0.122), followed by MLP (0.685 mean correlation for training versus 0.545 for test with a gap of 0.139). In the endothelial dataset, generalization was remarkably tight for the top sequence models (the transformer encoder achieved 0.500 training correlation versus 0.470 test correlation, with a gap of 0.031, and the LSTM achieved 0.462 training correlation and 0.453 test correlation, with a gap of 0.009). These small gaps support the stability of the learning setup and indicate that performance differences across deep model families are not primarily due to overfitting. By contrast, several classical ensemble methods exhibited huge train-test gaps, consistent with their ability to fit noise in high-dimensional sparse features. For example, Extra Trees reached almost perfect correlation (≈1.0) in both embryonic and endothelial cells. In contrast, the test correlation was much lower (0.442, with a gap of 0.558 in embryonic cells and 0.428 with a gap of 0.572) in endothelial cells. Similar patterns were seen for XGBoost (e.g., endothelial training correlation ≈ 1.0 vs test correlation 0.410). Together, these results suggest that model capacity alone is not sufficient: architectural bias and regularization behavior strongly shape whether capacity translates into out-of-sample accuracy under a fixed cis-feature representation.

**Fig. 4.**
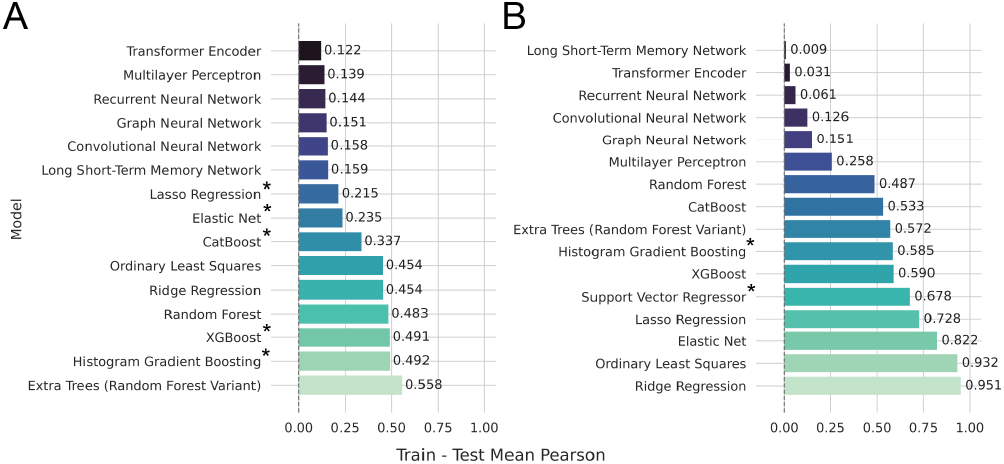
Generalization gaps across datasets. Train–test generalization gaps (training minus test mean Pearson correlation) for each model family in the (A) embryonic and (B) endothelial datasets. Smaller gaps indicate better generalization, whereas significant gaps suggest overfitting under the fixed promoter-centered representation.

### Transformer encoder predictions track held-out expression at scale

To visualize model behavior, we plotted predicted versus observed expression values on held-out test cells for the best-performing model in each dataset (transformer encoder; Figure 5). Pooling gene–cell observations across modeled genes shows that the model captures broad expression trends while preserving clear dataset-specific differences in dispersion.

**Fig. 5.**
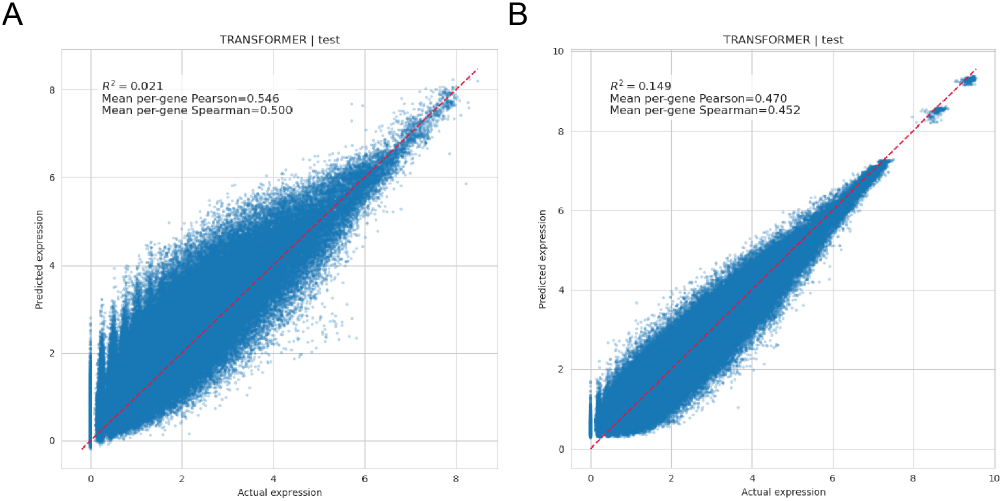
Predicted-versus-observed expression on held-out test cells. Scatter plots of predicted versus observed normalized expression values on the test set for the best-performing model (transformer encoder) in the (A) embryonic and (B) endothelial datasets. Each point corresponds to a gene–cell observation from the held-out test split (pooled across modeled genes). The diagonal line indicates perfect prediction (equality), illustrating dataset-dependent differences in dispersion and achievable accuracy under the promoter-centered feature representation.

Prediction behavior differed qualitatively across datasets, consistent with aggregate performance differences. In the embryonic dataset (transformer test mean 0.546), agreement between predicted and observed expression is tighter around the equality line. In contrast, in the endothelial dataset (test mean 0.470), scatter is more diffuse, consistent with weaker or less local chromatin-expression coupling and/or increased technical variability.

### Cis-regulatory feature importance is enriched near transcription start sites

We next examined which cisregulatory bins most strongly influenced predictions using SHAP-based feature importance (15) on the best-performing model (transformer encoder; Figure 6). Across both datasets, importance was enriched near the transcription start site and decayed with increasing distance, as highlighted in the zoomed panels. This pattern is consistent with the well-established role of promoter accessibility in transcription initiation (3, 4) and with the design of the SPEAR promoter-centered representation. Non-zero contributions from more distal bins within the window suggest that nearby non-promoter cis-regulatory activity still provides a proper predictive signal. Still, the sharp promoter-proximal enrichment indicates that much of the extractable signal in this feature space is concentrated close to the TSS, especially for genes whose expression is strongly accessibility-coupled.

**Fig. 6.**
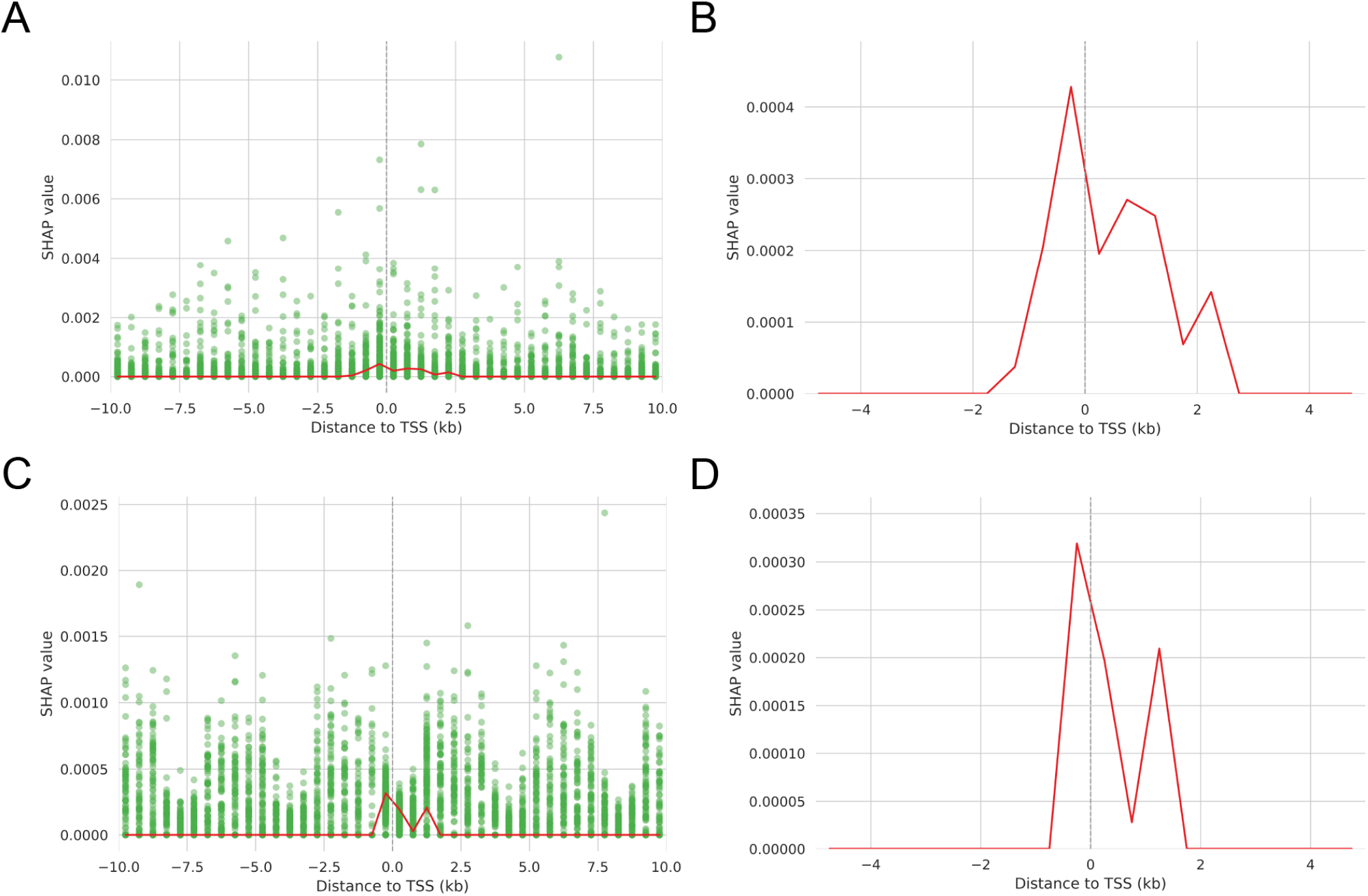
Cis-regulatory feature importance as a function of distance to the transcription start site (TSS). Relationship between cis-regulatory feature importance and genomic distance to the TSS, quantified using SHAP values from the best-performing model (transformer encoder). (A) Genome-wide feature-importance profile for the embryonic dataset. (B) Zoomed view of the embryonic dataset highlighting promoter-proximal regions. (C) Genome-wide profile for the endothelial dataset. (D) Zoomed view of the endothelial dataset highlighting promoter-proximal regions. Across datasets, importance is enriched near the TSS and decays with distance, consistent with promoter-proximal contributions dominating within the modeled window.

### Supplementary analyses reinforce robustness and model-dependent trade-offs

Supplementary analyses provide complementary views of performance and structure. Run-level summary statistics across models are reported in Supplementary Table 1. Alternative evaluation metrics (Spearman, R2, RMSE) broadly tracked the Pearson-based trends (Figure S2), while heatmap summaries emphasized gene- and model-specific structure (Figure S3). Gene-level extremes (Figure S4; transformer encoder only) underscore the long-tailed nature of predictability, showing that even the best model performs very well for some genes while remaining limited for others. Feature-level summaries (Figure S5; transformer encoder only) further show that a small number of bins dominate the model’s influence: the top cis-regulatory features by mean absolute SHAP value capture the most consistently informative genomic positions, reinforcing that the predictive signal in this formulation is sparse and position-dependent. Finally, the complexity– performance analysis (Figure S6) illustrates that increasing model complexity does not guarantee proportional gains, consistent with the observation that inductive bias (e.g., attention over ordered bins) is a major driver of accuracy under a fixed representation.

### Notes on fallback configurations

Models marked with an asterisk were evaluated using the 100-gene fallback manifest due to infeasibility or non-convergence at the 1,000-gene scale for the corresponding dataset. These results are included to document practical limitations of certain model classes in the high-dimensional (40 bins × genes) regime and to preserve comparability of the benchmarking study.

## Discussion

### Inductive bias dominates model performance under a fixed cis-regulatory representation

Across both datasets, deep neural architectures consistently outperformed classical baselines, with relative model rankings largely preserved despite substantial biological differences between systems. The transformer encoder achieved the highest mean test performance in both datasets (0.546 embryonic; 0.470 endothelial), indicating that attention-based architectures are particularly effective at extracting predictive structure from ordered cis-accessibility profiles. This stability in ranking suggests that model inductive bias, rather than dataset-specific idiosyncrasies, plays a dominant role under a fixed promoter-centered representation. Attention mechanisms provide a flexible way to integrate information across bins without enforcing strict locality or additivity, allowing transformers to capture distributed regulatory patterns within the ±10 kb window. While MLPs approached transformer performance in the embryonic dataset, structured sequence-aware models retained more evident advantages in the endothelial setting, consistent with the idea that architectural bias becomes more critical as signal-to-noise decreases or regulatory logic becomes more conditional. In contrast, classical linear models performed poorly, especially in the endothelial dataset, reflecting the difficulty of fitting high-dimensional, sparse, and correlated cis-accessibility features with purely additive assumptions. Tree-based ensembles occupied an intermediate regime, capturing nonlinearities but often suffering from overfitting in this setting.

### Biological context modulates absolute predictability

Although rankings were stable, absolute performance was consistently higher across the top models in the embryonic dataset. A plausible biological explanation is that early developmental programs exhibit tighter promoter–expression coupling, making promoter-centered accessibility particularly informative. In contrast, hemogenic endothelium likely relies more on distal enhancers, environmental responsiveness, and state-dependent regulation, reducing the fraction of expression variance explained by local accessibility alone. Technical factors likely contribute as well, including differences in sample size, sequencing depth, and sparsity profiles. Together, these observations suggest that promoter-centered cis accessibility provides a firm but context-dependent baseline for chromatin-to-expression prediction.

### Generalization behavior highlights the importance of architectural constraints

Split-wise evaluation and generalization gaps further emphasize the role of inductive bias. For the best-performing deep models, the train-test gap was modest, particularly in the endothelial dataset, indicating that improved performance reflects the extraction of genuine signal rather than memorization. In contrast, several ensemble methods exhibited extreme overfitting, achieving near-perfect training performance with substantially weaker test accuracy. These results underscore an essential point for chromatin-to-expression modeling: greater capacity alone does not guarantee better generalization. Architectural constraints that align with the structure of the feature representation, such as ordered bins and distributed cis-effects, appear more important than raw expressivity.

### Interpretability supports promoter-proximal dominance within the modeled regime

SHAP-based attribution revealed a strong enrichment of feature importance near transcription start sites in both datasets, with importance decaying with increasing distance from the transcription start site. This pattern aligns with established transcriptional biology (3, 4) and validates that the SPEAR representation captures biologically meaningful signals. While distal bins within the ±10 kb window contributed non-zero importance, the dominance of promoter-proximal bins suggests that much of the extractable predictive signal under this formulation is concentrated near the TSS. At the same time, the sparsity of influential features, highlighted by the top-ranked SHAP bins, suggests that relatively few positions disproportionately shape predictions, motivating future work on feature pruning and efficiency.

### Implications for transformer-based modeling of gene regulation

Transformer architectures have increasingly been applied to multimodal single-cell learning, including cross-modal prediction settings (e.g., multimodal transformer frameworks evaluated on ATAC-to-RNA tasks) (16). However, these approaches typically emphasize latent alignment and modality reconstruction, and are not designed to pro-vide controlled, gene-resolved benchmarking under a fixed cis-regulatory feature definition. The consistent performance of the transformer encoder in SPEAR suggests that attention-based models are particularly well-suited for this task. These results motivate broader adoption and deeper exploration of transformer-based architectures for chromatin-to-expression prediction, including variants that integrate distal regulatory information, motif-based trans features, or pretrained regulatory representations.

### Practical implications and limitations

From a practical perspective, SPEAR guides model selection under promoter-centered cis representations. When predictive accuracy is the primary objective, transformer encoders are a strong default. MLPs and regularized tree ensembles may be competitive in some contexts but require careful monitoring of generalization behavior. Several limitations frame these conclusions. SPEAR intentionally uses a fixed ±10 kb promoter-centered window, which likely underrepresents distal enhancer and trans-acting contributions. Default kNN smoothing improves stability but may inflate apparent predictability by borrowing information across similar cells. Finally, benchmarking was limited to two biological systems, and broader evaluation across tissues and perturbations will be required to establish generality.

### Future directions

SPEAR is designed to make targeted extensions straightforward. More broadly, accurate chromatin-to-expression prediction has practical implications for multimodal experimental design. When assays are typically limited to two (and less commonly three) modalities per cell, reliable prediction of RNA from ATAC could effectively free experimental capacity to profile additional regulatory layers in the same cells (17). Natural next steps include sweeping cis-window sizes to identify context-dependent regulatory scales, retraining models using only top-ranked features to quantify sparsity–performance trade-offs, and incorporating distal enhancer or transcription factor–related features to capture regulation beyond the promoter. Explicitly modeling cell type heterogeneity may further improve performance in complex tissues.

### Tool usability and reproducibility

Beyond benchmarking, SPEAR was designed as a practical, reproducible analysis platform. A configuration-driven workflow allows users to swap model families, adjust feature definitions, and modify preprocessing without altering code, while comprehensive run-level bookkeeping ensures reproducibility. Standardized outputs, including raw predictions, per-gene metrics, diagnostics, and interpretability analyses, enable down-stream reanalysis and extension without retraining. Together, these design choices position SPEAR as both a benchmarking framework and a foundation for future methodological development in single-cell regulatory modeling.

## Data availability

All datasets analyzed in this study are publicly available. The mouse embryonic multiome dataset is available from GEO under accession GSE205117. The human hemogenic endothelium multiome dataset is available from GEO under accession GSE270141.

## Code availability

SPEAR is open source and publicly available at https://github.com/UzunLab/SPEAR. The repository includes the complete Python implementation, command-line interface for training and inference, and documentation, as well as the analysis scripts and notebooks used to generate all results and figures reported in this study.

## Competing interests

The authors have no competing interests to disclose.

## Author contributions statement

T.W.A. conceived and implemented the SPEAR framework, performed data preprocessing, model development, benchmarking experiments, and analyses, and drafted the manuscript. Y.U. supervised the project, contributed to the study design and interpretation of results, and edited the manuscript. All authors read and approved the final manuscript.

## ACKNOWLEDGEMENTS

This work was supported by start-up funds from the Penn State University College of Medicine, awarded to Yasin Uzun, the principal investigator.

## Notes

### Competing Interest Statement

The authors have declared no competing interest.

## Bibliography

1. Fuchou Tang, Catalin Barbacioru, Yangzhou Wang, Ellen Nordman, Clarence Lee, Nanlan Xu, Xiaohui Wang, John Bodeau, Brian B. Tuch, Asim Siddiqui, Kaiqin Lao, and M. Azim Surani. mRNA-Seq whole-transcriptome analysis of a single cell. Nature Methods, 6(5):377–382, May 2009. ISSN 1548-7105. doi: 10.1038/nmeth.1315.

2. Saiful Islam, Una Kjällquist, Annalena Moliner, Pawel Zajac, Jian-Bing Fan, Peter Lönnerberg, and Sten Linnarsson. Characterization of the single-cell transcriptional landscape by highly multiplex RNA-seq. Genome Research, 21(7):1160–1167, July 2011. ISSN 1088-9051, 1549-5469. doi: 10.1101/gr.110882.110.

3. Jason D. Buenrostro, Beijing Wu, Ulrike M. Litzenburger, Dave Ruff, Michael L. Gonzales, Michael P. Snyder, Howard Y. Chang, and William J. Greenleaf. Single-cell chromatin accessibility reveals principles of regulatory variation. Nature, 523(7561):486–490, July 2015. ISSN 1476-4687. doi: 10.1038/nature14590.

4. Junyue Cao, Darren A. Cusanovich, Vijay Ramani, Delasa Aghamirzaie, Hannah A. Pliner, Andrew J. Hill, Riza M. Daza, Jose L. McFaline-Figueroa, Jonathan S. Packer, Lena Christiansen, Frank J. Steemers, Andrew C. Adey, Cole Trapnell, and Jay Shendure. Joint profiling of chromatin accessibility and gene expression in thousands of single cells. Science, 361(6409):1380–1385, September 2018. doi: 10.1126/science.aau0730.

5. Sai Ma, Bing Zhang, Lindsay M. LaFave, Andrew S. Earl, Zachary Chiang, Yan Hu, Jiarui Ding, Alison Brack, Vinay K. Kartha, Tristan Tay, Travis Law, Caleb Lareau, Ya-Chieh Hsu, Aviv Regev, and Jason D. Buenrostro. Chromatin Potential Identified by Shared Single-Cell Profiling of RNA and Chromatin. Cell, 183(4):1103–1116.e20, November 2020. ISSN 0092-8674, 1097-4172. doi: 10.1016/j.cell.2020.09.056.

6. Kevin E. Wu, Kathryn E. Yost, Howard Y. Chang, and James Zou. BABEL enables cross-modality translation between multiomic profiles at single-cell resolution. Proceedings of the National Academy of Sciences, 118(15):e2023070118, April 2021. doi: 10.1073/pnas.2023070118.

7. Tim Stuart, Andrew Butler, Paul Hoffman, Christoph Hafemeister, Efthymia Papalexi, William M. Mauck, Yuhan Hao, Marlon Stoeckius, Peter Smibert, and Rahul Satija. Comprehensive Integration of Single-Cell Data. Cell, 177(7):1888–1902.e21, June 2019. ISSN 0092-8674, 1097-4172. doi: 10.1016/j.cell.2019.05.031.

8. Yuhan Hao, Stephanie Hao, Erica Andersen-Nissen, William M. Mauck, Shiwei Zheng, Andrew Butler, Maddie J. Lee, Aaron J. Wilk, Charlotte Darby, Michael Zager, Paul Hoffman, Marlon Stoeckius, Efthymia Papalexi, Eleni P. Mimitou, Jaison Jain, Avi Srivastava, Tim Stuart, Lamar M. Fleming, Bertrand Yeung, Angela J. Rogers, Juliana M. McElrath, Catherine A. Blish, Raphael Gottardo, Peter Smibert, and Rahul Satija. Integrated analysis of multimodal single-cell data. Cell, 184(13):3573–3587.e29, June 2021. ISSN 0092-8674, 1097-4172. doi: 10.1016/j.cell.2021.04.048.

9. Jeffrey M. Granja, M. Ryan Corces, Sarah E. Pierce, S. Tansu Bagdatli, Hani Choudhry, Howard Y. Chang, and William J. Greenleaf. ArchR is a scalable software package for integrative single-cell chromatin accessibility analysis. Nature Genetics, 53(3):403–411, March 2021. ISSN 1546-1718. doi: 10.1038/s41588-021-00790-6.

10. Sneha Mitra, Rohan Malik, Wilfred Wong, Afsana Rahman, Alexander J. Hartemink, Yuri Pritykin, Kushal K. Dey, and Christina S. Leslie. Single-cell multi-ome regression models identify functional and disease-associated enhancers and enable chromatin potential analysis. Nature Genetics, 56(4):627–636, April 2024. ISSN 1546-1718. doi: 10.1038/s41588-024-01689-8.

11. Ricard Argelaguet, Tim Lohoff, Jingyu Gavin Li, Asif Nakhuda, Deborah Drage, Felix Krueger, Lars Velten, Stephen J. Clark, and Wolf Reik. Decoding gene regulation in the mouse embryo using single-cell multi-omics, June 2022. Pages: 2022.06.15.496239 Section: New Results.

12. Harmke Biezeman, Martina Nubiè, and Leal Oburoglu. Hematopoietic cells emerging from hemogenic endothelium exhibit lineage-specific oxidative stress responses. Journal of Biological Chemistry, 300(11), November 2024. ISSN 0021-9258, 1083-351X. doi: 10.1016/j.jbc.2024.107815.

13. Tamara Goldfarb, Vamsi K Kodali, Shashikant Pujar, Vyacheslav Brover, Barbara Robbertse, Catherine M Farrell, Dong-Ha Oh, Alexander Astashyn, Olga Ermolaeva, Diana Haddad, Wratko Hlavina, Jinna Hoffman, John D Jackson, Vinita S Joardar, David Kristensen, Patrick Masterson, Kelly M McGarvey, Richard McVeigh, Eyal Mozes, Michael R Murphy, Susan S Schafer, Alexander Souvorov, Brett Spurrier, Pooja K Strope, Hanzhen Sun, Anjana R Vatsan, Craig Wallin, David Webb, J Rodney Brister, Eneida Hatcher, Avi Kimchi, William Klimke, Aron Marchler-Bauer, Kim D Pruitt, Françoise Thibaud-Nissen, and Terence D Murphy. NCBI RefSeq: reference sequence standards through 25 years of curation and annotation. Nucleic Acids Research, 53(D1):D243–D257, January 2025. ISSN 1362-4962. doi: 10.1093/nar/gkae1038.

14. Jonathan M Mudge, Sílvia Carbonell-Sala, Mark Diekhans, Jose Gonzalez Martinez, Toby Hunt, Irwin Jungreis, Jane E Loveland, Carme Arnan, If Barnes, Ruth Bennett, Andrew Berry, Alexandra Bignell, Daniel Cerdán-Vélez, Kelly Cochran, Lucas T Cortés, Claire Davidson, Sarah Donaldson, Cagatay Dursun, Reham Fatima, Matthew Hardy, Prajna Hebbar, Zoe Hollis, Benjamin T James, Yunzhe Jiang, Rory Johnson, Gazaldeep Kaur, Mike Kay, Riley J Mangan, Miguel Maquedano, Laura Martínez Gómez, Nourhen Mathlouthi, Ryan Merritt, Pengyu Ni, Emilio Palumbo, Tamara Perteghella, Fernando Pozo, Shriya Raj, Cristina Sisu, Emily Steed, Dulika Sumathipala, Marie-Marthe Suner, Barbara Uszczynska-Ratajczak, Elizabeth Wass, Yucheng T Yang, Dingyao Zhang, Robert D Finn, Mark Gerstein, Roderic Guigó, Tim J P Hubbard, Manolis Kellis, Anshul Kundaje, Benedict Paten, Michael L Tress, Ewan Birney, Fergal J Martin, and Adam Frankish. GENCODE 2025: reference gene annotation for human and mouse. Nucleic Acids Research, 53(D1):D966–D975, January 2025. ISSN 1362-4962. doi: 10.1093/nar/gkae1078.

15. Scott Lundberg and Su-In Lee. A Unified Approach to Interpreting Model Predictions, November 2017. 1705.07874 [cs].

16. Wenzhuo Tang, Hongzhi Wen, Renming Liu, Jiayuan Ding, Wei Jin, Yuying Xie, Hui Liu, and Jiliang Tang. Single-Cell Multimodal Prediction via Transformers. In Proceedings of the 32nd ACM International Conference on Information and Knowledge Management, pages 2422–2431, October 2023. doi: 10.1145/3583780.3615061. 2303.00233 [q-bio].

17. Chunyuan Yang, Yan Jin, and Yuxin Yin. Integration of single-cell transcriptome and chromatin accessibility and its application on tumor investigation. Life Medicine, 3(2), April 2024. doi: 10.1093/lifemedi/lnae015.

