## Supplemental Information for "SPEAR: Predicting Gene Expression from Single-Cell Chromatin Accessibility"

**This PDF file includes:**
Supplementary Text
Figs. S1 to S6
Table S1 legend (full table provided separately)

### Supplementary Text

This supplementary document compiles additional benchmarking analyses for SPEAR, including split-wise performance summaries, expanded metric distributions, heatmap-based comparisons, gene-level predictability extremes, SHAP-based feature rankings, and model-complexity comparisons. Together, these figures provide additional views of model performance, robustness, interpretability, and complexity under the fixed promoter-centered representation used throughout the manuscript. The legend for Supplementary Table 1 is included at the end of this PDF, while the full table is provided separately.

### Supplementary Figures

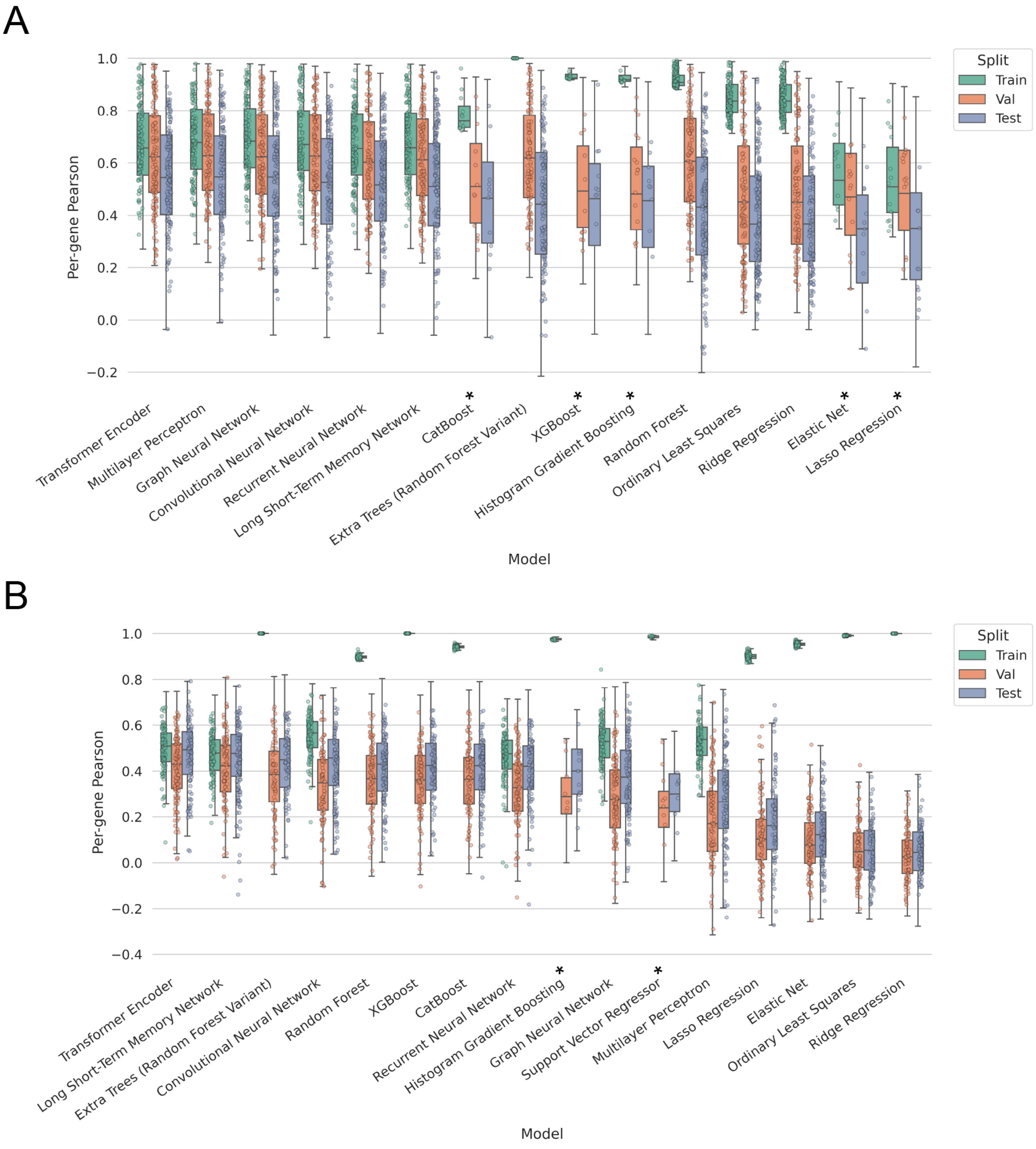


**Fig. S1. Split-wise performance consistency across datasets.** Mean Pearson correlation performance across training, validation, and test splits for each model family in the (A) embryonic and (B) endothelial datasets. Consistent split-wise performance indicates stable generalization, while divergences highlight potential overfitting to high-dimensional promoter-centered features.

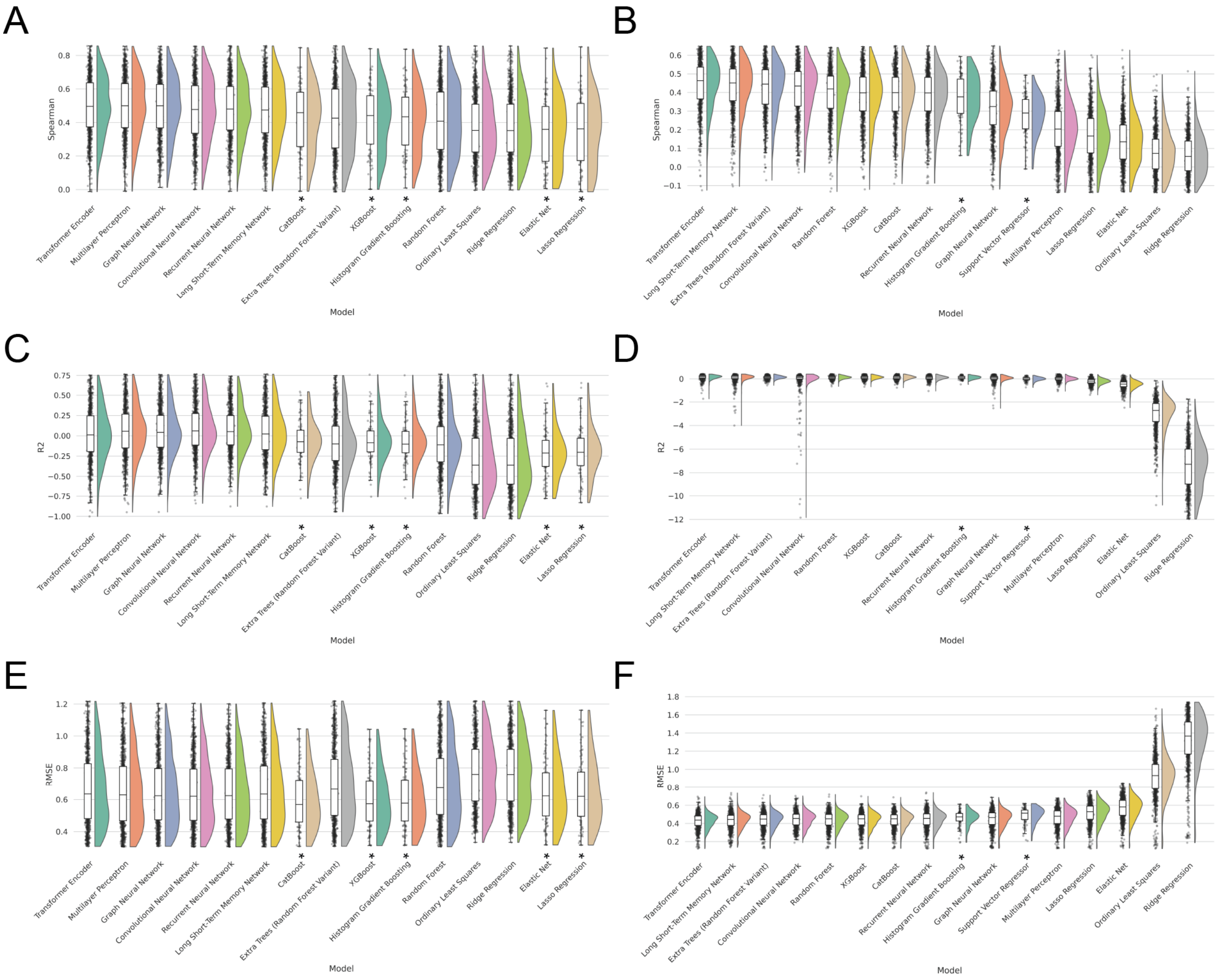


**Fig. S2. Expanded performance distributions across evaluation metrics.** Distributions of gene-level test performance across model families for complementary metrics: (A) embryonic Spearman correlation, (B) endothelial Spearman correlation, (C) embryonic $R^2$, (D) endothelial $R^2$, (E) embryonic RMSE, and (F) endothelial RMSE. Together, these metrics provide additional views of ranking stability and error structure beyond Pearson correlation. Models evaluated using the 100-gene fallback manifest are indicated with an asterisk.

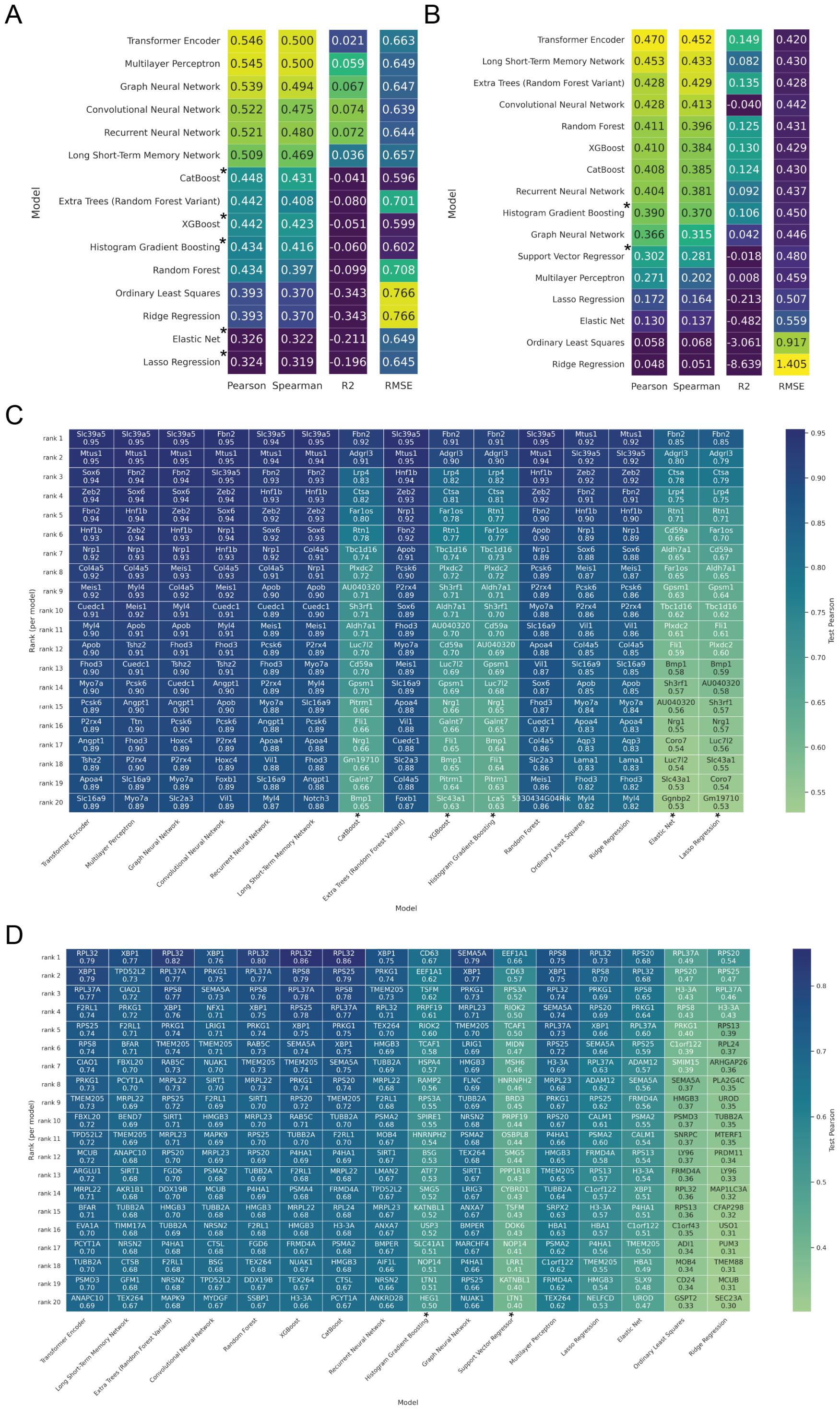


**Fig. S3. Heatmap-based summaries of model performance.** (A–B) Heatmaps of mean test-set performance across model families for the embryonic (A) and endothelial (B) datasets (metrics as shown). (C–D) For each model family, heatmaps of the top-performing genes by test Pearson correlation in the embryonic (C) and endothelial (D) datasets; entries report gene identity and corresponding test correlation, highlighting gene- and model-specific structure in predictability.

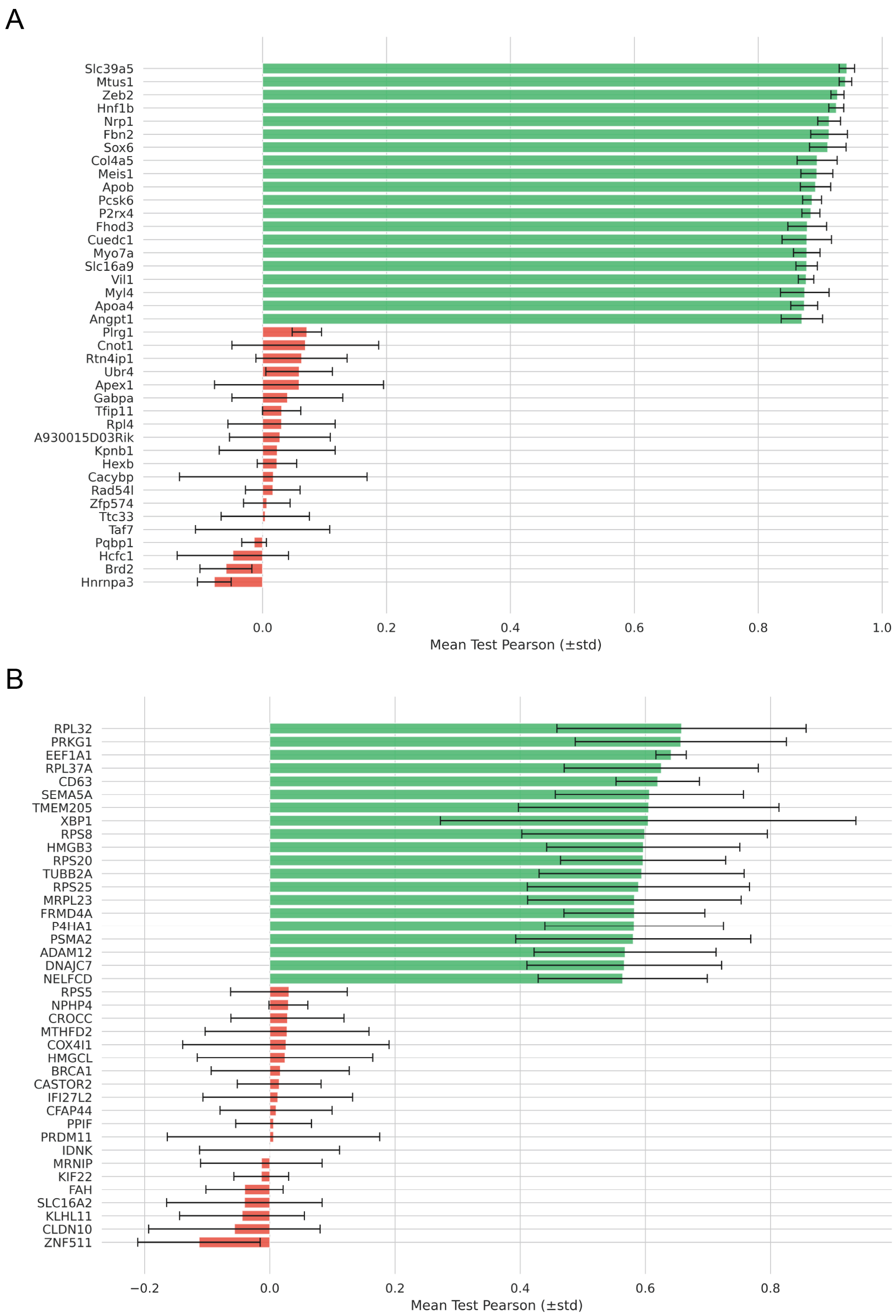


**Fig. S4. Extremes of gene-level predictability.** Top and bottom genes ranked by mean test-set Pearson correlation under the best-performing model (transformer encoder) in the (A) embryonic and (B) endothelial datasets. Error bars indicate the reported variability measure (as plotted), illustrating the long-tailed nature of gene predictability under promoter-centered accessibility features.

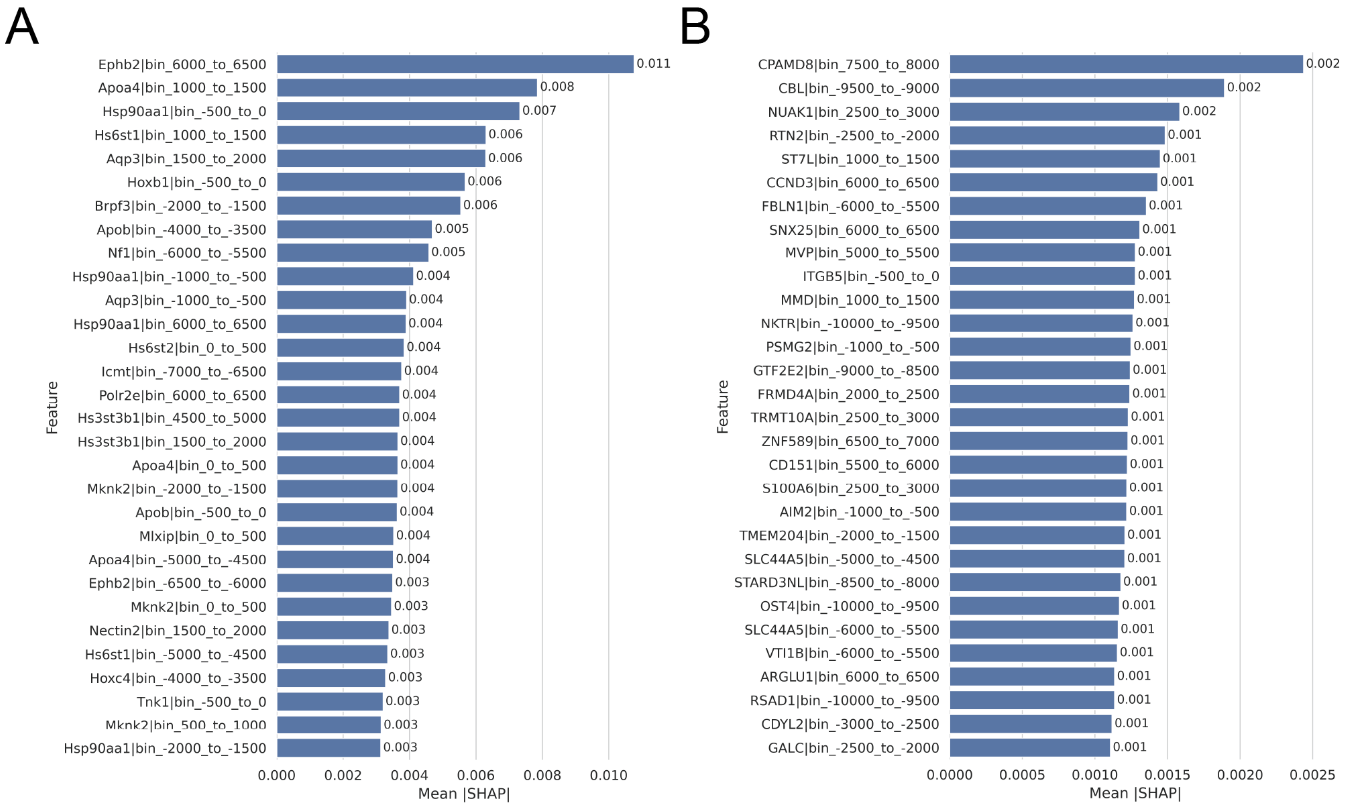


**Fig. S5. Top cis-regulatory features ranked by absolute SHAP importance.** Top 30 cis-regulatory bins ranked by mean absolute SHAP value for the transformer encoder in the (A) embryonic and (B) endothelial datasets. Absolute SHAP values quantify the magnitude of contribution to predictions irrespective of direction, highlighting genomic positions with the strongest and most consistent influence on predicted expression.

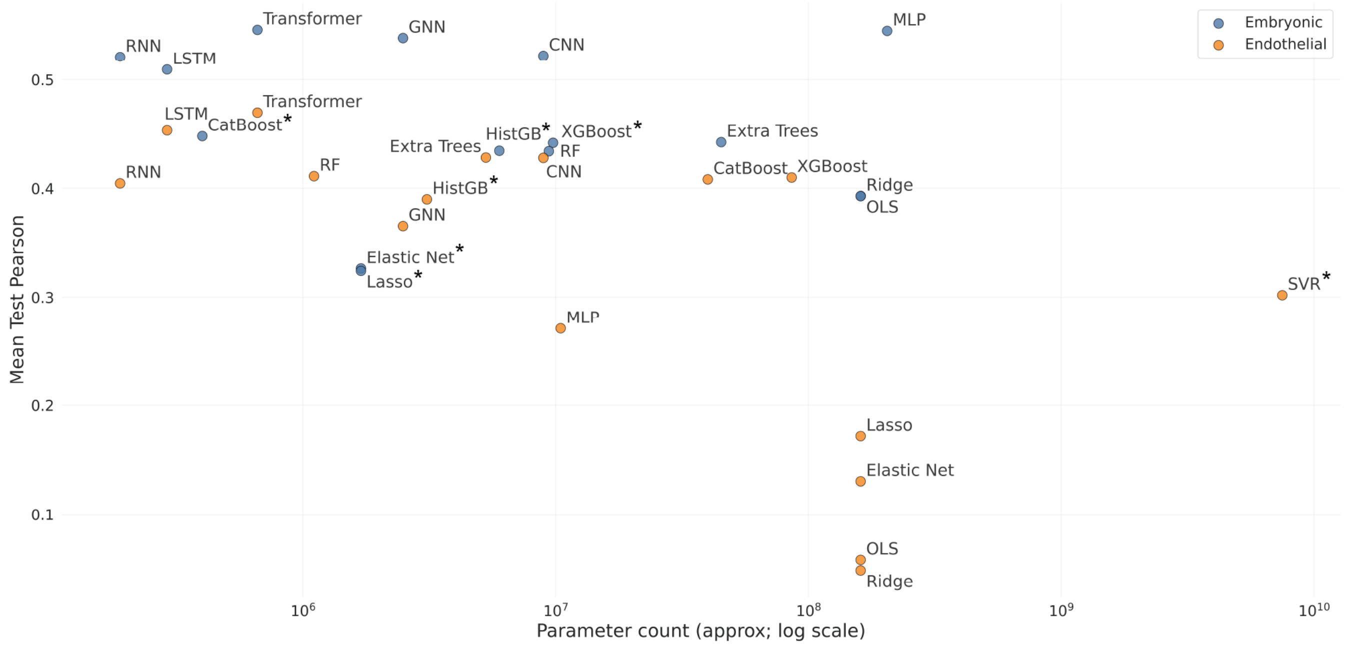


**Fig. S6. Relationship between model complexity and predictive performance.** Mean test-set Pearson correlation as a function of model complexity (parameter count; plotted on a log scale) across model families and datasets. The analysis highlights performance–complexity trade-offs under a fixed promoter-centered representation and shows that increased complexity does not necessarily yield proportional gains.

### Supplementary Tables

**Supplementary Table 1. Run-level summary statistics across evaluated model families.** Summary metrics for each model–dataset run in the mouse embryonic development and human hemogenic endothelium benchmarks. The attached table reports the display name, dataset, internal model identifier, and run name for each benchmarked configuration, together with mean and standard deviation across modeled genes for training, validation, and test Pearson correlation, Spearman correlation, $R^2$, RMSE, MSE, and MAE. Models marked with an asterisk in the manuscript were evaluated using the 100-gene fallback manifest because the corresponding 1,000-gene configuration was infeasible or did not converge reliably.
